# Vapor mediation as a tool to control micro-nano scale dendritic crystallization and preferential bacterial distribution in drying respiratory droplets

**DOI:** 10.1101/2021.06.18.448992

**Authors:** Omkar Hegde, Ritika Chatterjee, Abdur Rasheed, Dipshikha Chakravortty, Saptarshi Basu

## Abstract

Deposits of biofluid droplets on surfaces (such as respiratory droplets formed during an expiratory event fallen on surfaces) are composed of the water-based salt-protein solution that may also contain an infection (bacterial/viral).The final patterns of the deposit formed are dictated by the composition of the fluid and flow dynamics within the droplet. This work reports the spatio-temporal, topological regulation of deposits of respiratory fluid droplets and control of motility of bacteria by tweaking flow inside droplets using non-contact vapor-mediated interactions. When evaporated on a glass surface, respiratory droplets form haphazard multiscale dendritic, cruciform-shaped precipitates—using vapor mediation as a tool to control these deposits at the level of nano-micro-millimeter scales. Wemorphologically control dendrite orientation, size and subsequently suppress cruciform-shaped crystals. The nucleation sites are controlled via preferential transfer of solutes in the droplets; thus, achieving control over crystal occurrence and growth dynamics. The active living matter like bacteria is also preferentially segregated with controlled motility without attenuation of its viability and pathogenesis. For the first time, we have experimentally presented a proof-of-concept to control the motion of live active matter like bacteria in a near non-intrusive manner. The methodology can have ramifications in biomedical applications like disease detection, controlling bacterial motility, and bacterial segregation.

## 1. Introduction

Drying patterns formed from the evaporation of droplets of complex biological fluids(such as tears, synovial fluid or spinal fluid, blood, etc.) are enigmatic and have implications in biomedical applications[1–3] such as preventing disease transmission[4,5] and diagnostics[6,7].Bio-fluid droplets are complex fluids containing several constituents such as proteins,surfactants, salts, to mention a few[8]. Due to the many components present in the bio-fluid droplet, the competition between capillary flows driven by continuity and solutal Marangoni flows driven by surface tension gradients (due to differential evaporation of the components) determines the fluid flowinside a droplet[9]. Researchers have observed suppression of the coffee ring effect due to protein adsorption on the surface of the particles in protein droplets containing suspended polystyrene particles[10]. However, the deposit will be at the edge or uniformly distributed over the surface depending on the charge of the protein[11]. The presence of salts in bio-fluids further adds to the complexity leading to the formation of dendritic crystals. The dendritic structure formation is dependent on the salt concentration, drying mode, and particle size and shapes[12,13]. It is clear from several studies[13–16] that the final precipitate formed is dependent on the individual components, the ratio of the components present in the biofluid droplet, the substrate on which the droplet is evaporated, and the environmental conditions. Thus, the final morphology of a deposit depends on the combined effect of several parameters, as discussed, most importantly, mass transport due to fluid flow and the aggregation of colloidal particles within the droplet if parameters such as environmental conditions, substrate, components of the biofluid are maintained constant.

The flow inside droplets driven by evaporation is very low (~*O*(10)*μm/s*) and is highly uncontrolled. Since the flow inside the droplet is primarily responsible for the final deposit formed, the final deposit can be controlled by controlling the flow as a corollary. However, most techniques used to control the flow inside droplets, such as acoustic excitation[17], heating[18], magnetic stirring[19], the addition of surfactant[20], are highly intrusive and can lead to denaturing of the biological sample. Hence we propose the non-intrusive vapor mediated interaction[21,22] to control the flow and subsequent patterns formed on drying of the bio-fluid droplet. This is done by placing an ethanol droplet in the vicinity of the bio-fluid droplet (we use a pendent ethanol droplet as shown in Figure 1 (b)), thus creating an asymmetric concentration field of ethanol around the biofluid droplet. The minuscule amount of ethanol vapor in the vicinity of the bio-fluid droplet is adsorbed onto its surface, creating a surface tension gradient across the bio-fluid droplet. This generates vigorous Marangoni flow inside the bio-fluid droplet, whose magnitude and flow direction can be controlled by strategically positioning the ethanol droplet in the vicinity of the bio-fluid droplet (as shown in Figure 1 (b), (c), and (d)).

**Figure 1.**
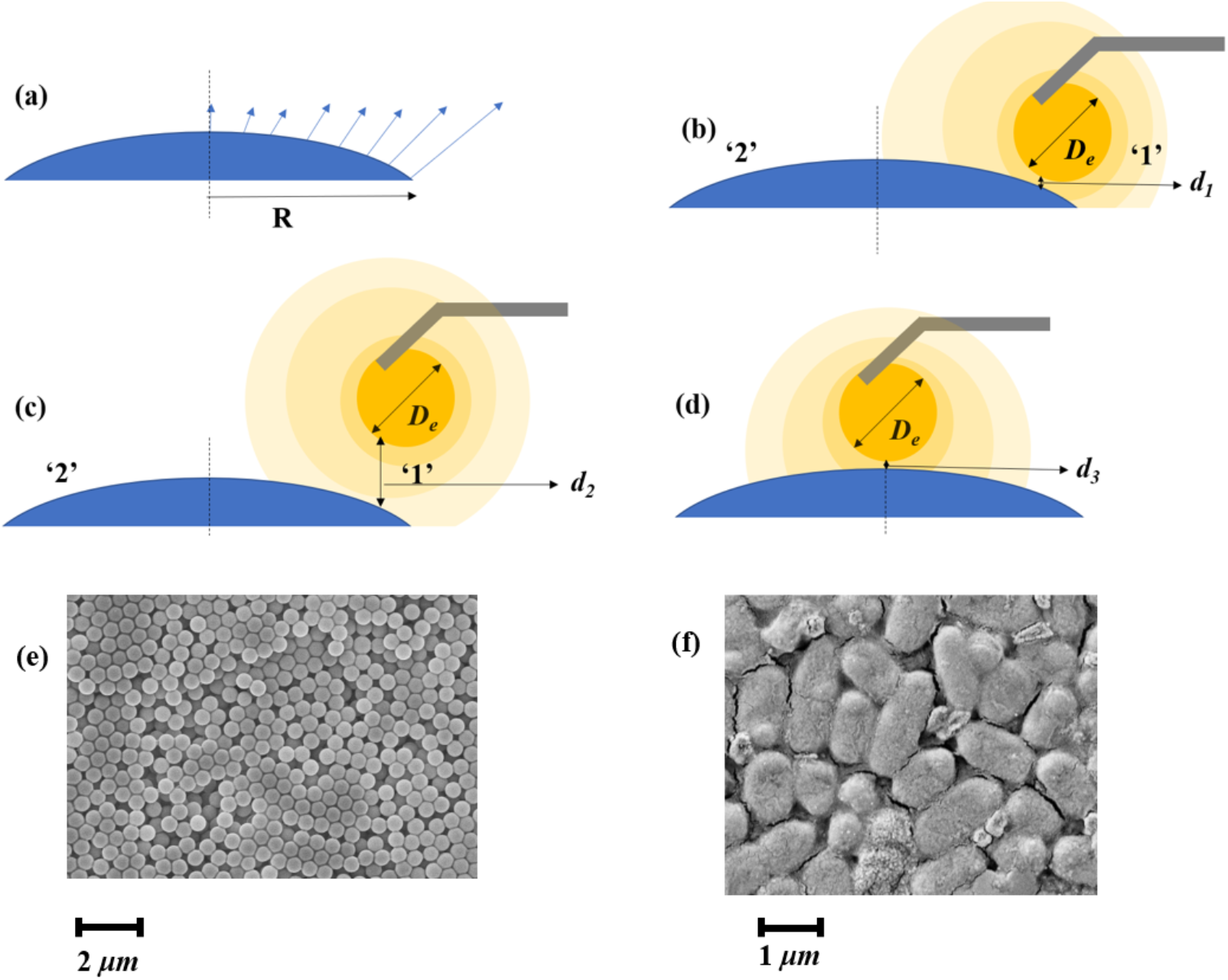
Schematic representation of the experimental cases. The Surrogate Fluid Droplet (SRF) is allowed to evaporate on the glass surface in the following configurations (a) case 1: A single SRF droplet is placed, (b) case 2: a pendent ethanol droplet is brought very close to the SRF at a distance of *d_1_*~0.085 ± 0.01 *mm* on the side ‘1’, (c) case 3: a pendent ethanol droplet is placed at a farther distance (*d_2_*~0.21 ± 0.03 *mm*) to the SRF droplet on side ‘1’, (d) case 4: a pendent ethanol droplet is placed very close to the SRF droplet at a distance *d_3_*~0.085 ± 0.025 *mm*at the center of the drop. SEM image of (e) 860 *nm* inert polystyrene microspheres, (f) rod-shaped STM bacteria.

Besides several components present in the bio-fluids, bacteria are also present in the bio-fluid droplets and are generally motile[23], unlike the inert micro/nanosuspension in the droplets[9,10]. Bacterial motility in these droplets is key to its navagation[24], biofilm formation[25], and self-assembly[26] and is crucial for its survival in the dried precipitates[27]. Bacteria also respond to the change in the environment[28] and the surfaces in their vicinity[29]. Experimental observations in the literature indicate that the bacteria exhibit swimming motion in droplets that can even propel the droplets[30]. Several methods have been devised to engineer the bacterial motion[31], which can potentially be used to develop biosensors[32] and other microfluidic devices[33]. However, microbial motility with respect to its physical environment, most notably to fluid flow in its surroundings, is often neglected in the literature. In order to see the effect of flow on the deposition and aggregation of live bacteria in the bio-fluid droplets, we seed rod-shaped *Salmonella enterica* serovar Typhimurium as a model system in the bio-fluid. Although *Salmonella* Typhimurium (STM), a gut pathogen, is transmitted by the orofecal transmission route via contaminated food and water, many studies have demonstrated the aerosol transmission of specific serovars of *Salmonella enterica*, such as *Salmonella* Typhimurium, *Salmonella* Agona, etc. [34,35]. With this as a physiological signifance, STM is used in the experiments as a model system. In the present work, surrogate respiratory fluid (SRF) (for components, see section 2.1) droplets containing bacteria (*Salmonella enterica* serovar Typhimurium) are considered a model bio-fluid system to demonstrate the control of crystalization and bacterial deposition through non-intrusive vapor mediation. In a practical scenario, respiratory droplets generated during an expiratory event from an infected host may fall on the ground, can form fomites, and have a potential for secondary disease transmission[5]. In this article, vapor mediation is used as a tool for preferential bacterial deposition on the evaporated deposit without diminishing its viability and simultaneously retaining its pathogenesis.In addition, we report the transformation ofspatio-temporal and topological regulation of crystals formed in bio-fluid droplets via controlled Marangoni convection. Multiscale dendritic cruciform-shaped precipitates are formed on the drying of a single surrogate respiratory droplet (without the presence of vapor). We have demonstrated that thedendrite orientation and size can be morphologically controlled, and subsequently, we can suppress cruciform-shaped crystals using vapor mediation. The nucleation sites are also controlled via preferential transfer of solutes in the droplets; thus, achieving control over occurrence and crystal growth dynamically.

## 2. Experimental Section

### 2.1 Preparation of Surrogate Respiratory fluid (SRF)

The surrogate model of respiratory fluid used in experiments consists of dissolved salts and alveolar surfactants emulating reality[7]. The respiratory fluid composition used in this article is the same as Vejerano et al. reported[36]. The process of preparation of the surrogate fluid is detailed as follows: 0.9 % by wt. of NaCl, 0.3% by wt. of gastric mucin (Type III, Sigma Aldrich), and 0.05 % wt. of di-palmitoyl phosphatidyl-choline (DPPC (Avanti Polar Lipids)) is added in deionized water. The final composition is sonicated for 15 minutes to create a homogeneous solution. Next, the homogenized solution is centrifuged at 5000 RPM for 15 minutes to pellet the impurities present in the liquid. The pH value of the prepared solution is 6 or greater.

### 2.2 Preparation of bacterial culture

Fluorescently labeled mCherry wild type (WT) *Salmonella enterica* serovar Typhimurium strain 14028 (S. Typhimurium) is grown in Luria broth with appropriate antibiotic concentrations under shaking conditions at 37°C. Overnight cultures prepared from a single colony from a freshly streaked plate are used for the experiments.1.5*ml* of overnight culture was centrifuged at 6,000 RPM to pellet the bacterial cells and washed once with autoclaved MilliQ water. The resulting pellet is then resuspended into a freshly prepared surrogate respiratory solution (SRF) and serially diluted such that each SRF droplet of 0.5 *μl*approximately contains ~10^3^ bacteria. The approximate size and shape of the bacteria used are shown in Figure 1(f).

### 2.3 Experimental set-up

0.5±0.1*μl* droplet of surrogate respiratory fluid (SRF) (with bacteria) is gently placed on the clean glass substrate as shown in Figure 1 (a) and allowed to evaporate in controlled laboratory conditions (temperature 27± 3°C, and relative humidity at 40±5 %). This is referred to as case 1 in this article. Next, a 2 *μl* pendent ethanol droplet is brought near side ‘1’ (see Figure 1(b) and (c)) of SRF droplet at a distance of *d_1_*=0.085 ± 0.01 *mm* and *d_2_*=0.21± 0.03 *mm* referred to as case 2 and case 3, respectively (refer to Figure 1 (b) and (c), Side ‘1’ is referred to the side where the ethanol is placed and side ‘2’ is the side opposite to the ethanol). Case 4 consists of the 2 *μl* pendent ethanol droplet being place close to the SRF droplet near the center of the SRF and the distance between them being *d_3_*=0.085 ± 0.025 *mm* (refer to Fig. 1 (d)). The distances mentioned above (*d_1_*, *d_2_*, *d_3_*) are the distance between the pendant ethanol droplet’s surface and the surface of the SRF droplet, which is maintained constant until the SRF droplet evaporates (as shown in Figure 1). The pendent ethanol droplet volume is maintained constant throughout the experiment by maintaining a constant pumping rate of 1 *μl/ minute* (for the given laboratory conditions) equivalent to ethanol evaporation.

All experiments are conducted at least four times to maintain repeatability and reproducibility.

The contact line dynamics and the droplet’s height are obtained from shadowgraphy images using a NIKON D7200 camera attached to a Navitar zoom lens. The volume of the droplet is estimated, assuming the droplet to be of a spherical cap. The crystallization dynamics are captured at 2.19 *fps* from the top-view, imaged using a high-resolution CCD camera (PCO2000) mounted on a BX51 Olympus frame (See Video 7.1, Video 7.2, Video 7.3, Video 7.4). A halogen-based light source (TH4 200, Olympus) is used for top-view illumination. *μ*-PIV experiments are done to study the flow field within the droplet qualitatively and qualitatively. Neutrally buoyant monodisperse polystyrene particles are used for *μ*-PIV experiments,as shown in Figure 1 (e) (the same was used in our previous work[37]). The settings for *μ*-PIV experiments are the same as described in our previous work[37].

### 2.4 Viability and Infection Assay

To assess the viability of the bacteria in the dried droplets, the precipitate is resuspended in 40*μl* of PBS and plated onto *Salmonella-Shigella* agar (SS agar) at appropriate dilution. The viability of bacteria is calculated by multiplying with dilution factor in terms of CFU/*ml*. Further, to measure the pathogenicity of these bacteria in dried droplets, the resuspended droplets are subjected tomurine macrophages RAW 264.7.Further, 24 well plates are centrifuged at 500-700 RPM to enhance bacterial attachment to host cells and incubated for 25minutes at 37°C and 5% CO_2_. The media containing bacteria is discarded and washed thrice with 1X phosphate buffer saline (PBS).The cells are further subjected to Gentamicin treatment (dissolved in DMEM) at a concentration of 100 *μg/mlfor* 1 hour to eliminate any extracellular bacteria. The cells are maintained at 25*μg/ml* gentamicin containing DMEM for the entire experiment. Finally, infected cells are lysed using 0.1% Triton-X 100 at 2hours and 16 hours post-infection, and appropriate dilutions spread on SS agar plates.

The fold proliferation is calculated as follows: CFU at 16 hours is divided by CFU at 2 hours to obtain fold replication of intracellular bacteria.

## 3. Results and Discussions

### 3.1 Spatio-temporal control of crystallization

In this section, we describe the global dynamics of crystallization as obtained from the experimental observations. In the SRF, 70% of the solute mass is NaCl. On drying of NaCl solution, cuboidal crystals are expected to form[37]. However, due to the presence of other colloids (mucin, surfactant), a gelatinous mixture is formed at the later stages of evaporation due to solvent desiccation. The salt solution is non-homogeneously dispersed and is embedded in the gelatinous matrix.The formation of dendrites is due to the non-homogenous distribution of salt in the droplet. The role of mucin in the present study is analogous to agar forming dendrites, as observed by Goto et al.[38].The inception of crystallization occurs when the solution attains supersaturated conditions due to the drying of the droplet. For Case 1, the solute is accumulated more near the edge of the droplet; thus, the rim of the droplet gets saturated faster. The optical profilometry data of the dried droplet shows a ring deposit with a ring thickness of ~3-4 *μm*(refer to FigureS6 in supplementary information). As a result of supersaturation near the rim of the droplet, the onset of crystallization always occurs from the droplet’s rim, which acts as a nucleus, and thereby, there is sustained growth of crystals connected to the saturated solution that propagates to the center[9] (Refer to Video 1 in the supplementary information, Figure 2 (a)).Crystallization can start anywhere from the rim (as it is an instability), and there is no control over it with the natural evaporation of the droplets. The instantaneous length of the dendrite (l) is considered from the point of nucleation until the tip of the growing front.

**Figure 2.**
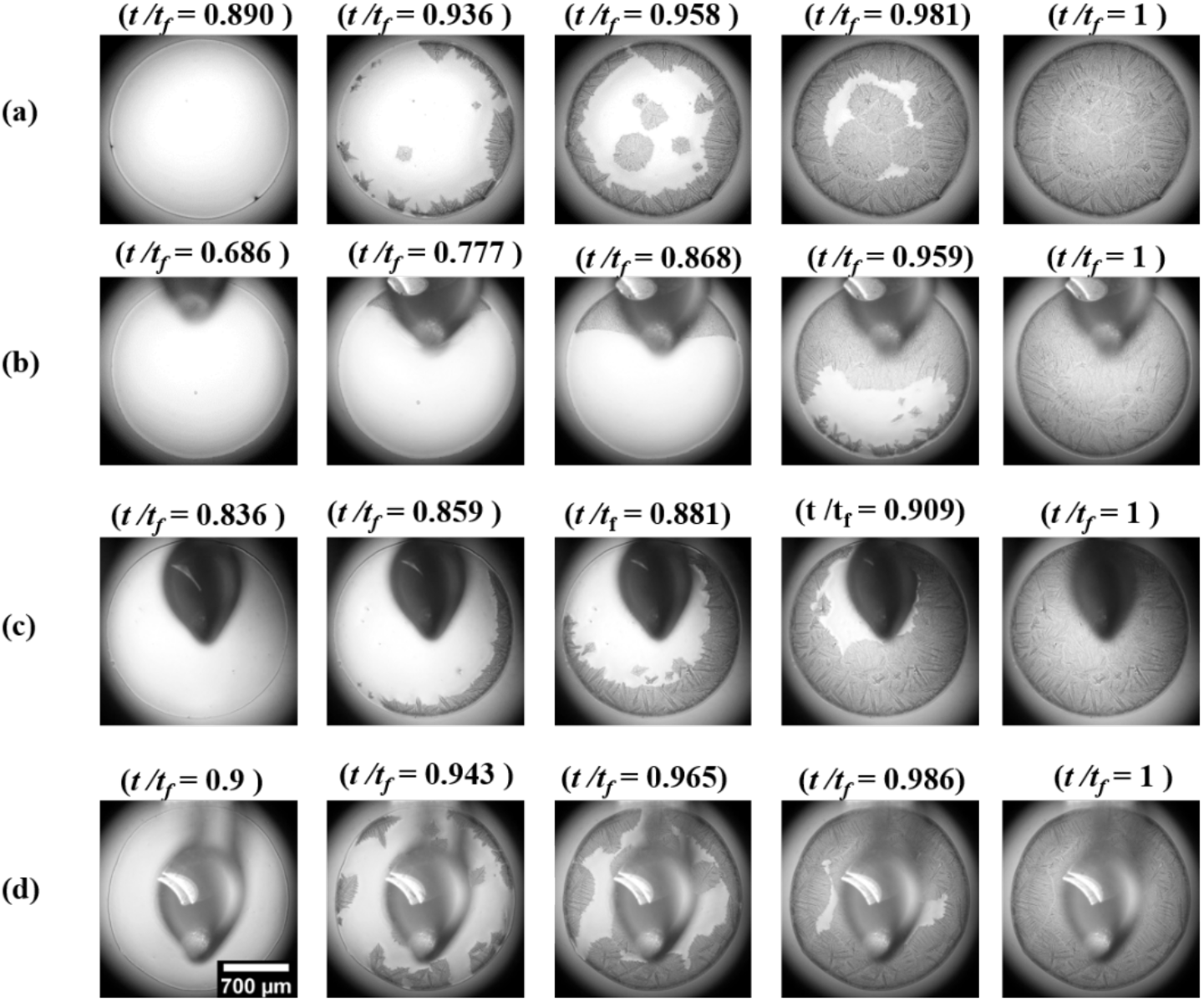
Dynamics of crystalization. Snapshots with non-dimensional time *t/t_f_* (given at top of every image) for (a) case 1, (b) case 2, (c) case 3, (d) case 4. The scale bar given in the bottom left image in (d) is valid for all images in the Figure. *t* is the instantaneous time, and *t_f_* is the total time of evaporation.

We now show that by changing the flow inside the droplet through vapor mediation, we can preferentially segregate the solute within the droplet, thus control the inception of crystalization. A pendent ethanol droplet brought close to the SRF droplet at a distance *d_1_* (case 2, refer to Figure 1 (b)). Due to the proximity of the ethanol droplet, there is vigorous Marangoni flow induced in the sessile droplet[22],which leads to a contact line slip from side ‘1’ (see Video2, Figure 2 (b)) (flow dynamics inside the droplet are further described in detail in section2.3, and we will currently focus on the global contact line and crystallization dynamics in this section). The contact line slip was observed in salt solution droplets in our previous work[37]. However, in the present work, although the slip occurs from the side ‘1’, it leaves behind a gelatinous substance at the side ‘1’ near the initial contact line due to colloids present in the SRF droplet. As the fluid entirely moves towards side ‘2’, leaving behind trace solute near side ‘1’, the region near side ‘1’ attains saturation. Thus the inception of crystalization for case 2 always happens from side ‘1’ (Video 7.2, Figure 2 (b)). However, since the contact line motion is due to liquid flow towards side ‘2’ and not as a consequence of evaporation, there is a considerable amount of liquid present in the side ‘2’. Hence, although the inception of crystalization occurs very fast, the growth of crystalization is slow compared to other cases (refer to Figure S1 in the supplementary information). Due to the contact line slip, an uneven ring deposit with a lesser thickness ~ 1.7 *μm* at side ‘1’ and more thickness ~ 5 *μm* at side ‘2’ is formed in case 2 (refer to Figure S7 the supplementary information).

The strength of the Marangoni flow is reduced when the ethanol droplet is placed at a distance *d_2_*(as*d_2_*>*d_1_*, see section 2.3), which is insufficient to cause contact line slip. However, more solute is deposited on the side ‘2’ on every flow circulation due to Marangoni flow. This is a similar type of deposition observed in our previous work[21]. As a result, the inception of crystalization is reversed (starts from side ‘2’) due to supersaturation at side ‘2’ of the droplet (refer to Figure 2 (c) and Video 3). Thus, although there is no contact line slip, there is a preferential transfer of solute to side ‘2’. This is reflected in the optical profilometry data with a thicker deposit at side ‘2’ (refer to Figure S8 in the supplementary information).

When the ethanol droplet is brought near the center of the sessile droplet (case 4, Figure 1 (d)), the crystalization occurs from the center as well as the rim of the droplet, merging to join each other (refer to Figure 2 (d) and Video 3). As an experimental constraint, it is not possible to observe crystalization from the center of the droplet due to the hindering pendent ethanol droplet (refer to Figure 2 (d)). Hence the time for the inception of crystalization is taken as the time when we first see the crystals from the droplet’s rim (Figure S1(b) in the supplementary information). As the fluid and solute flow outwards from the center, the central region of the dried precipitate has the most negligible thickness, and the thickness of the deposit increases radially (refer to Figure S9 in the supplementary information).

### 3.2 Multiscale dendritic patterns

SEM Micrographs reveal that the dendritic crystals on the dried SRF droplet are of size ranging from ~*O*(1) *μm* to ~*O*(10^3^) *μm*. The crystal pattern form in a haphazard way from randomly formed nucleation sites for case 1. Crystals grow from the inner nucleation site in cruciform dendritic shape with branches radiating outwards(refer to Figure S2 in the supplementary information).

The formation and growth of dendritic crystals depend on competition between solvent loss (which increases the concentration of salt) and diffusion of salt ions towards the leading edge of the tip (which reduces the concentration of salt in solution)[9]. With the natural evaporation of the droplet, there is no control over the length, shape, or dynamics of crystallization. As a result, crystals of different shapes are formed everywhere on the deposit.

It has been observed from the experiments that a graded distribution of crystal sizes and controlled directional orientation can be obtained by controlling the flow in droplets using vapor mediation. When the ethanol droplet is placed very close to the SRF droplet (case 1), fine tiny crystals of ~*O*(1) *μm* to ~*O*(10) *μm* are observed in the slip region at side ‘1’ (refer to Figure 3 (d)). Suppression of cruciform-shaped crystals is observed near side ‘1’. With the crystalization starting from side ‘1’, elongated dendrites greater than ~*O*(10^2^) *μm* with their orientation away from the side ‘1’ are formed (refer to Figure 3 (c)). Certain crystals form at an inclined orientation from side ‘1’ (refer to Figure 3 (b), (e), and (f), which could be due to a non-controlled vapor source in the very vicinity, leading to vigorous Marangoni in different directions. A controlled vapor field, as done by Volpe et al.[39] could reduce such aberrations, which is out of the scope of the present work as we are dealing with only the proof-of-the-concept. Since the region near side ‘2’ is less affected by the vapor field, we see the dendritic crystals are similar to case 1, such as the haphazard formation of cruciform crystals (refer to Figure S3 in supplementary information).

**Figure 3.**
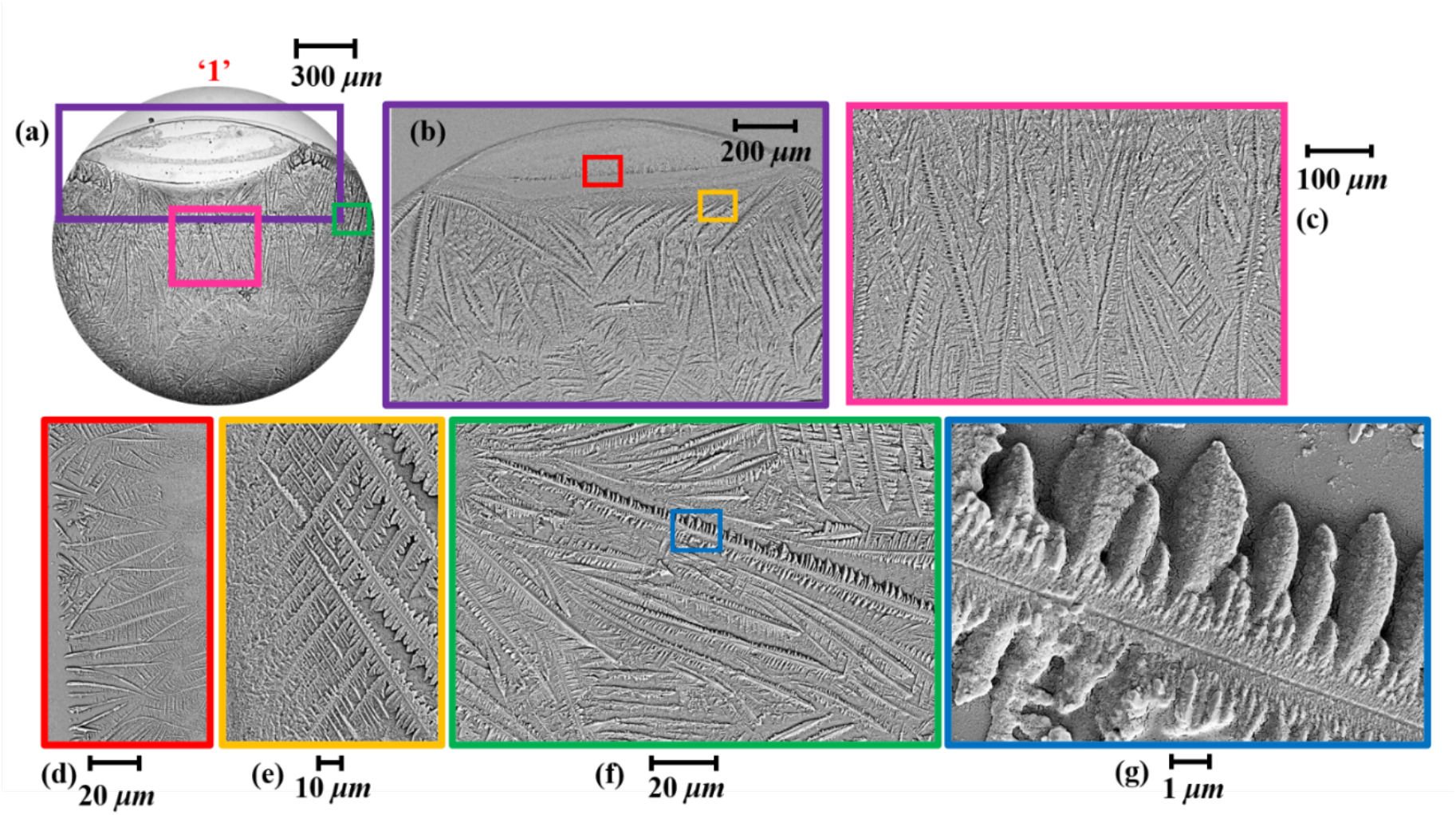
SEM images of dried precipitate for case 2 near side ‘1’ (colored). (a) Image of the deposit near side ‘1’. Zoomed-in image of the region within the box, (b) with purple border in (a), (c) with pink border in (a), (d) with a red border in (b), (e) with a yellow border in (b), (f) with green border in (a), (g) with a blue border in (f).

We observe that the cruciform-shaped crystals are suppressed near side ‘1’ even for case 3. The directional flow from side ‘1’ to ‘2’ in the droplet creates elongated parallel dendrites with directional orientation from side ‘1’ to side ‘2’ (refer to Figure S4 in the supplementary information). Similar to case 2, cruciform-shaped crystals are found near side ‘2’ for case 3. In case 4, a central hole is formed at the droplet in the end stages of evaporation, similar to a hole formed in thin films due to vapor-mediated Marangoni convection[40]. We again report a graded size distribution of the dendrites for case 4, with the smallest being near the center (~2-5 *μm*) and bigger dendrites (~100-300 *μm*) as we move radially outwards. The dendrites are oriented towards the radial direction (to Figure S5 in the supplementary information).

### 3.3 Flow dynamics inside droplet under the influence of vapor mediation

The flow inside the droplet is measured using μ-PIV. The flow inside an SRF droplet (Case 1 in this chapter) was analyzed by Abdur et al.[9]. Flow is visualized in the SRF droplet by seeding fluorescent particles of 860 *nm*procured from Thermo Fischer (refer to Video 6). All measurements are taken below the midplane of the droplet (*h/h_0_*<0.5). Double toroidal Marangoni flow is observed initially in the SRF droplet, which later transforms into the capillary flow near the evaporation. The magnitude of flow is ~*O*(10) *μm/s(refer* to Figure4 (a)). The flow in case 2 is initially circulatory and becomes unidirectionally away from side ‘1’ later (refer to Figure4 (b)). At later times the surface tension gradient is very high, and due to the unidirectional flow, the contact line slips. The explanation for this was given in our previous studies[37].

**Figure4.**
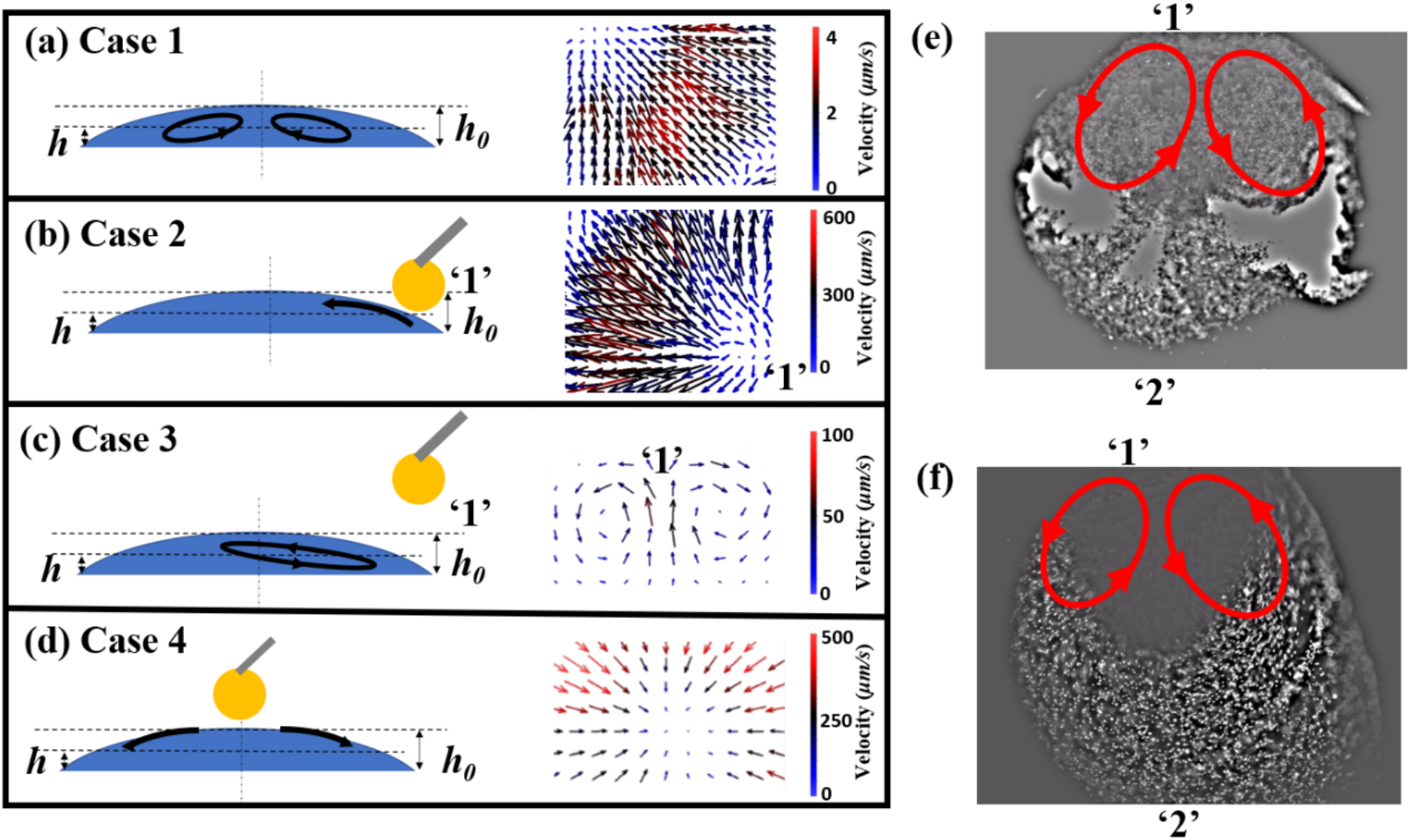
Instantaneous vectors of velocity for (measurement *h/h_0_*<0.5taken below the mid-plane of the droplet (a) case 1 (at initial times ~*t/t_f_*~0.1), (b) case 2 (‘1’ indicates the side at which the ethanol is placed, measurement at ~*t/t_f_*>0.7), (c) case 3 (‘1’ indicates the side at which the ethanol is placed, measurement at ~*t/t_f_*>0.7), (d) case 4 (at initial times ~*t/t_f_*~0.1). (e) Flow visualization for case 2 with bacteria (refer to video 5). Bacteria tend to aggregate near side ‘2’. (f) Flow visualization for case 2 with 860 *nm* inert nanoparticles (refer to video 6) where the particles do not aggregate (are discrete). The red arrows in (e) and (f) represent the flow direction.

The magnitude of flow velocity in case 2 is found to be ~*O*(10^3^) *μm/s*. In case 3, the Marangoni convection is insufficient to cause a slip, but circulatory flow with a lower magnitude of flow ~*O*(10^3^) *μm/s*occurs. There is unidirectional flow at the end of evaporation in case 3, similar to case 2; however, the flow is insufficient to cause a slip. The flow inside SRF droplet in case 2 and case 3 is qualitatively similar; only the magnitude of velocity differs by one order of magnitude. In case 4, there is a circulatory flow to maintain the continuity; however, as the droplet thickness becomes significantly less in the end, strong radially outward flow from the center creates a dent at the center. The mechanism of formation of holes was previously explained by Kim et al.[40] is similar to case 4.

### 3.4 Bacterial distribution on the dried deposit

The inertial forces generated by the fluid flow in the droplet (~*O*(10^3^) *μm/s*)are very high compared to the bacterial motility. Therefore, it is expected that the bacteria move along with the flow as it would be difficult for them to resist inertia due to the Marangoni flow. At the end stage of evaporation, a gelatinous matrix of colloids formed is highly viscous, and the bacteria are trapped in it. Therefore, the motility of the bacteria will largely be restricted once the droplet dries. With the proposition that the bacteria faithfully follows the flow inside the droplets, we expect that most of the bacteria must lie in the regions of maximum solute deposit (information of the solute deposit in terms of thickness of the deposit is obtained from optical profilometry data).The SEM images do not reveal the final bacterial positions as the bacteria might have been embedded in the crystal matrix. However, since the bacteria used are tagged with a red fluorescent protein, confocal microscopy reveals the final positions of the bacteria in the deposit. Confocal microscopy data shows that most bacteria are found at the midplane of the crystal deposit.

Figure 5 (a) shows that the intensity is maximum near the ring because of the high bacterial density in the region. Since most of the solute deposit in case 1 is at the ring, bacteria are also found near the ring. The ring deposit is a consequence of the fluid flow, as discussed in section 2.3. Ring deposit due to solutal Marangoni flow was also explained by Marin et al.[41].

**Figure 5.**
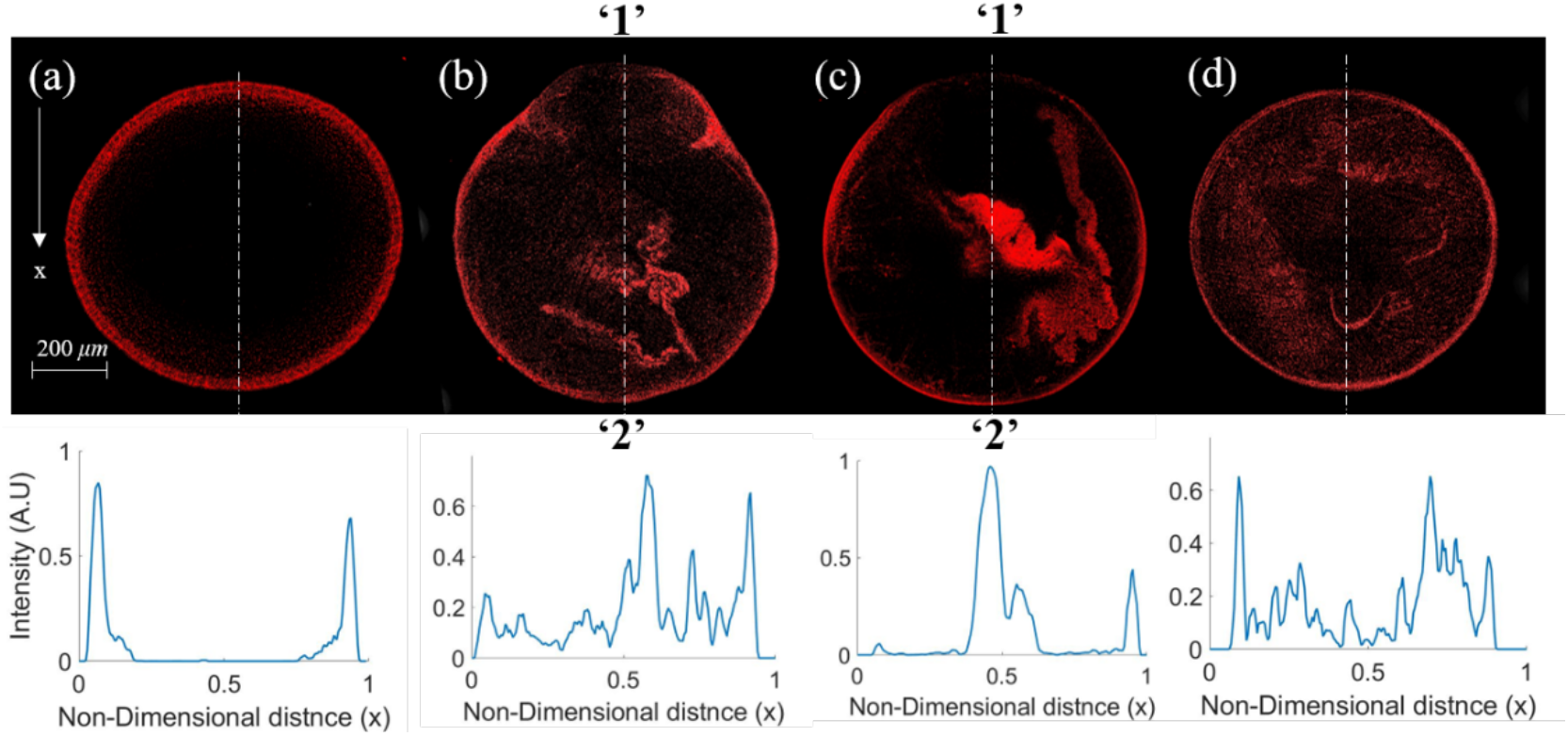
Confocal microscopy of the deposit with the fluorescence emission from the bacteria, (a) case 1, (b) case 2, (c) case 3, (d) case 4. Plots in the bottom row correspond to the intensity variation along the dashed line in the top row.

The bacterial deposit for cases 2 and 3 is less toward side ‘1’ compared to side ‘2’ (Figure 5 (b) and (c)). The bacterial deposit is less near the center for case 4 (Figure 5 (d)).

The fluorescent polystyrene particles used in μ-PIV experiments (which are inert and faithfully follow the flow as they are neutrally buoyant and have low stokes number of the particles) show similar deposits as bacteria (refer to Figure S10 in the supplementary information). However, clustered agglomeration is seen in the case of bacteria. Such agglomeration of bacteria could be due to the bacterial response to high shear flow[42]. The clustering of bacteria is seen by fluorescence visualization of the bacteria in the Marangoni flow (see Figure 4 (f) and refer to video 6). Thus the bacteria do not just move along with the flow. However, the confocal images in Figure 5 show that bacterial distribution can be controlled on a global scale using vapor mediation.

### 3.5 Bacterial distribution and viability of the dried precipitate

We have described in our previous works that the process of vapor mediation is non-intrusive as a minimal amount of ethanol is adsorbed on the surface of the droplet[21,43]. This can be seen from the volume regression curves of the SRF droplet (refer to Figure S11 in the supplementary information). The evaporation characteristics of the SRF droplet remain unchanged irrespective of the presence of ethanol droplet in its vicinity for all the four cases described (refer to Figure S11 in the supplementary information). This section investigates if there is any effect of vapor mediation on bacterial viability and its pathogenicity.

Bacterial reminiscent in the deposit survives over hours and days[44]. To see the effect of vapor mediation on the deposit, we conduct viability tests as described in section 2.3. The deposit after two hours is plated, and viable bacteria count is noted. Case 1 is considered as a control. The viability count in case 1 is found to be ~ 5×10^4^ CFU/*ml*. Percent viability is calculated by non-dimensionalizing the viability count of a particular case to that of Case 1. Viability for case 2, case3, and case 4 remain the same as case 1, as shown in Figure 6 (a). Thus, the bacterial viability is invariant of ethanol vapor present in the vicinity of the SRF droplet. However, when an ethanol droplet of 0.5 *μl* is directly cast onto the dried deposit of case 1, bacteria loses its viability. This case where ethanol comes in direct contact is labeled as case x. Thus it is clear that vapor mediation is non-intrusive to the viability of the bacteria. The uptake of bacteria by murine macrophages RAW264.7 (phagocytic cell) is observed, and the percentage of phagocytosis is calculated. The percentage of phagocytosis is the same for all cases (refer to Figure6 (b), case 1 is taken as a control, and values of other cases are non-dimensionalized by case 1). The viable bacteria have the potential to infect the host through the oro-fecal route. The bacteria, which has now been internalized, will be able to replicate inside the host cell. The fold proliferation data reveals the survival and virulence of the bacteria after entering the host cells. Fold proliferation of all four cases is retained at the same level (refer to Figure 6 (c)).

**Figure 6.**
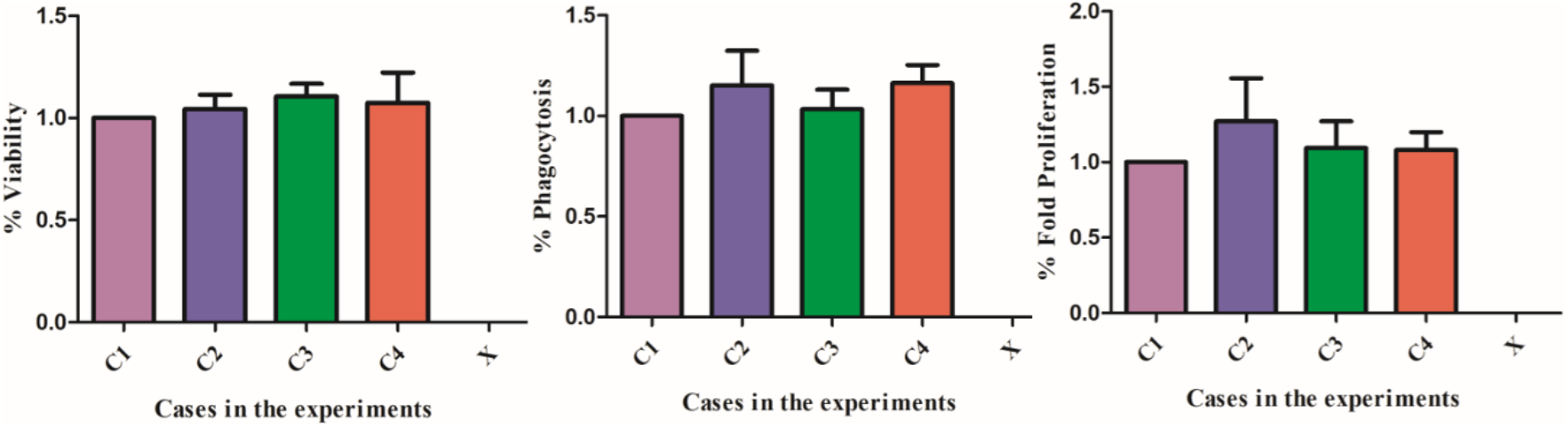
Plots of (a) percentage viability, (b) percentage phagocytosis, (c) percentage fold proliferation of bacteria for different cases. Case x represents when 0.5 *μl* of (volume of ethanol droplet is equivalent SRF droplet deposited) of 99% ethanol droplet is poured onto the deposit and allowed to dry.

A new culture is prepared (4 times each), and the bacteria resuspended in the SRF is drop-casted onto the surface, and drying experiments are conducted for all four configurations of Case 1,2,3 and 4 with four trials each. Thus, considering multiple freshly prepared cultures and multiple droplets cast onto the surface of each freshly prepared culture for all configurations give us the same results. Therefore, the error bar plotted in Figure 6 is calculated, taking into account the trials mentioned above.

## 4. Conclusions

Manipulation or segregation of particles in drying droplets[45], controlling motion of droplets using vapor mediated interaction is shown by several studies[21,39,46–48]; however, controlling an active living matter such as bacteria using a non-intrusive vapor mediation technique has not been studied. We have experimentally investigated the control of the distribution of bacteria within the deposit using vapor mediation without affecting bacterial viability and pathogenesis. The flow inside the SRF droplet is altered by strategic positioning of ethanol droplet leading to spatio-topological control of self-assembly, organization of biomolecules, and crystallization is demonstrated using vapor mediated interaction of droplets. Multiscale dendritic patterns can be formed and dynamically controlled using this technique. This study provides valuable insights into the vapor-mediated non-contact mechanism to control thepattern formation in complex solutions like biofluids. These findings provide a preliminary understanding of the bacterial interaction with the fluid flow inside the droplet. Proof of concept presented in this work can be used as a tool to control bacterial motility, its segregation, and patterns of bio-fluid deposits which may have wide-ranging implications in clinical infection scenario and biomedical engineering.

## Notes

### Competing Interest Statement

The authors have declared no competing interest.

