## Supplementary Materials for "Vapor mediation as a tool to control micro-nano scale dendritic crystallization and preferential bacterial distribution in drying respiratory droplets"

**Supplementary information:**

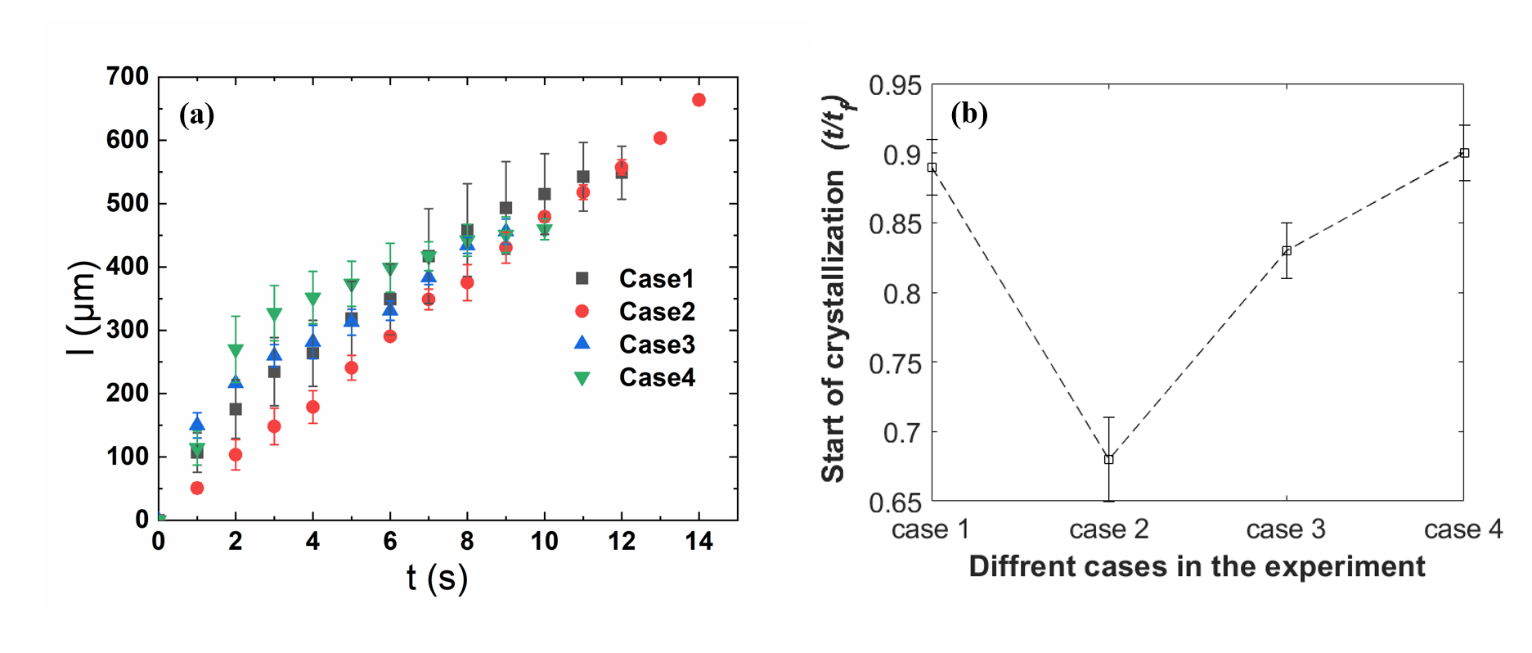

Figure S1 Growth dynamics of the crystal. (a)The plot of instantaneous dendrite length (l) with time (t). (b) the plot of the approximate time of inception of crystalization for different cases.

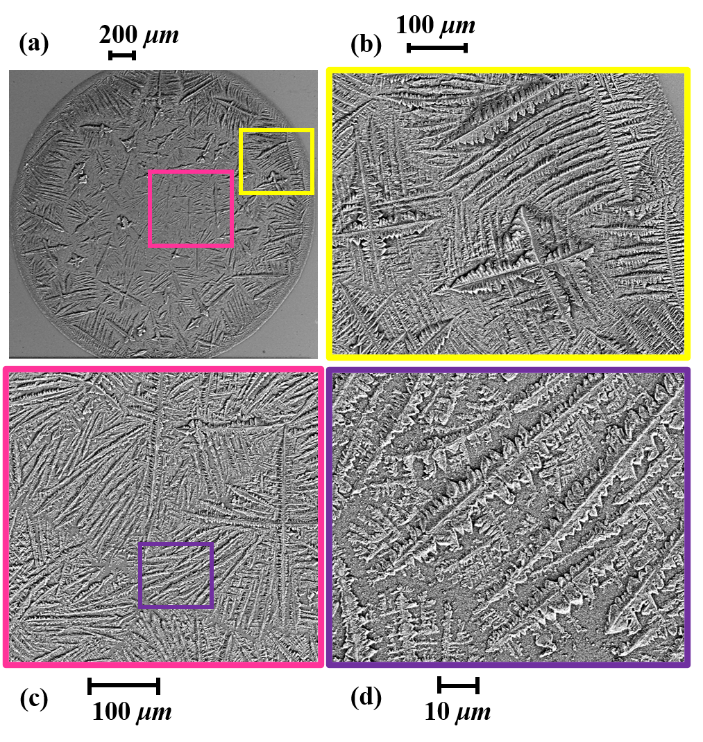

Figure S2 SEM images of dried precipitate for case 1 (colored). (a) Image of the entire deposit. Zoomed-in image of the region within the box, (b) with a yellow border in (a), (c) with pink border in (a), (d) with yellow purple in (c).

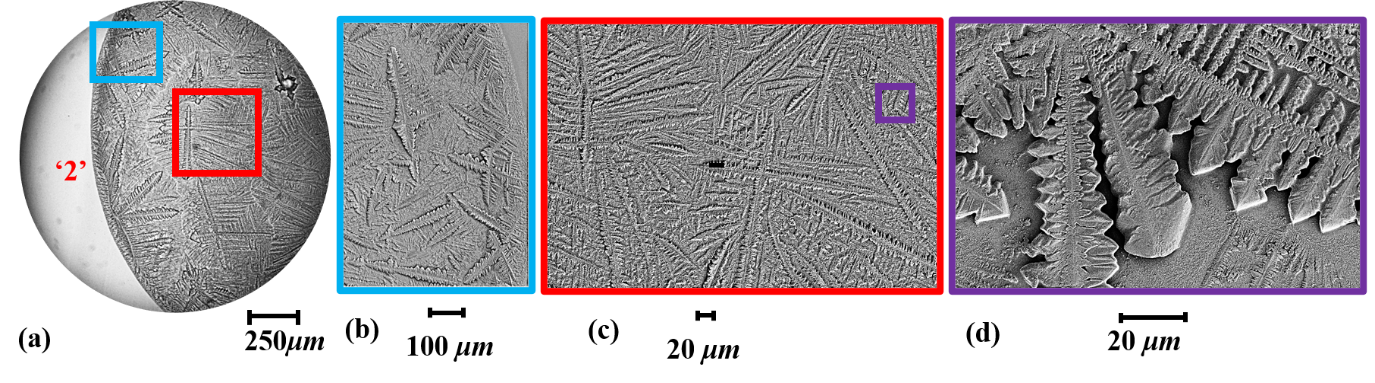

Figure S3 SEM images of dried precipitate for case 2 near side ‘2’ (colored). (a) Image of the deposit near side ‘2’. Zoomed in image of, (b) the region within the box with sky blue border in (a), (c) the region in the box with a red border in (a), (d) the region within the box with a purple border in (c).

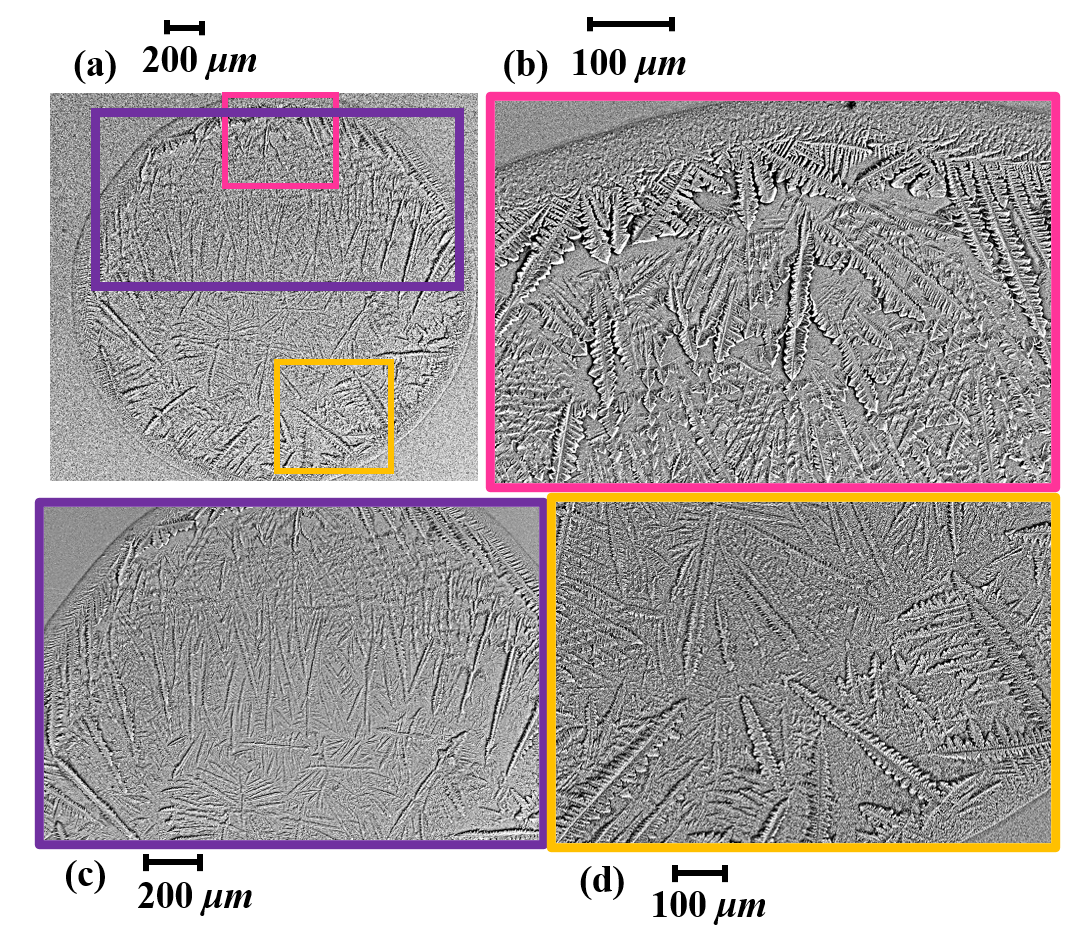

Figure S4 SEM images of dried precipitate for case 3 (colored). (a) Image of the entire deposit. Zoomed-in image of the region within the box, (b) with pink border in (a) which is near side ‘1’, (c) with purple border in (a), (d) the box with a yellow border in (a) which is near side ‘2’.

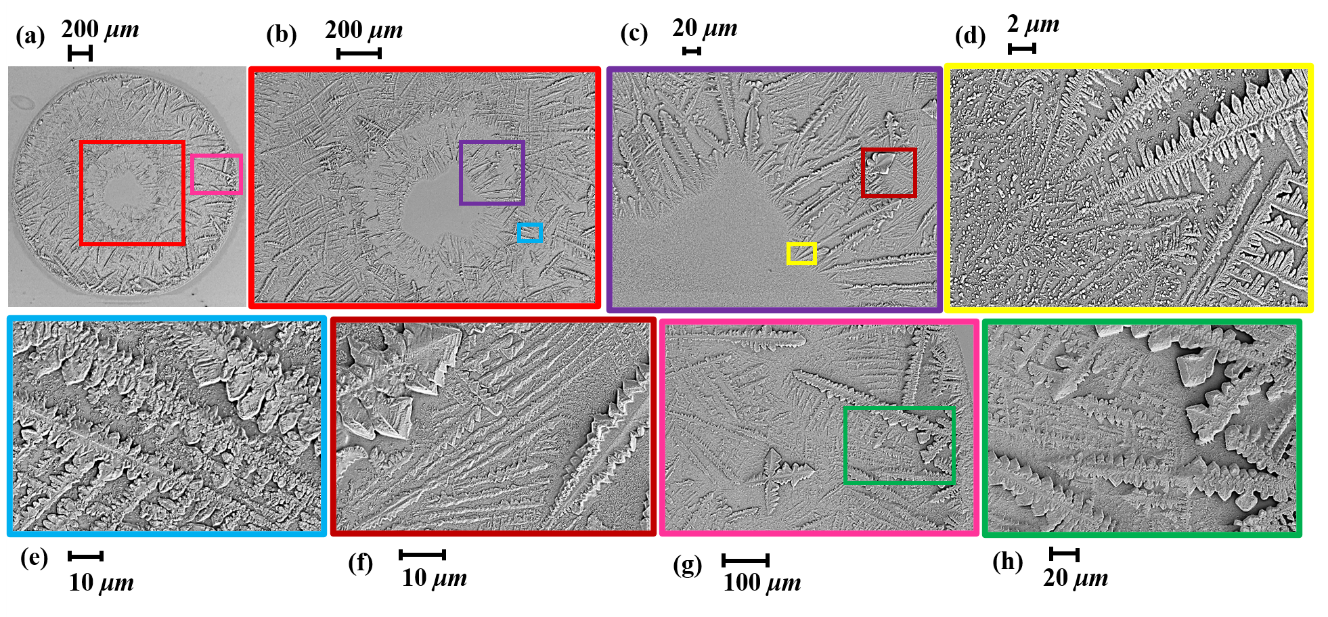

Figure S5 SEM images of dried precipitate for case 4 (colored). (a) Image of the entire deposit. Zoomed-in image of the region within the box, (b) with a red border which is at the center of (a), (c) with purple border in (b), (d) with a yellow border in (c), (e) with a sky blue border in (b), (f) with a brown border in (c), (g) with a brown border in (c), (h) with a brown border in (c).

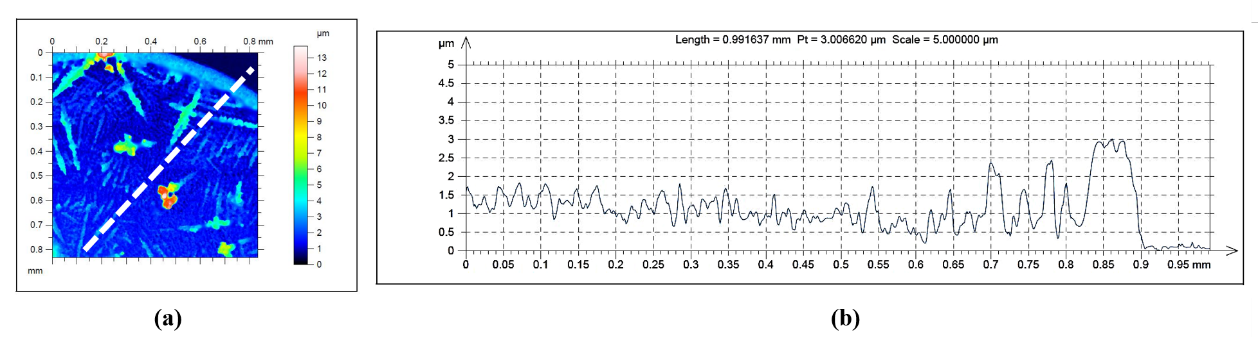

Figure S6 Optical profilometry data of dried precipitate for case 1, (a) 2D surface profile, (b) Variation of the height along the dashed line in (a).

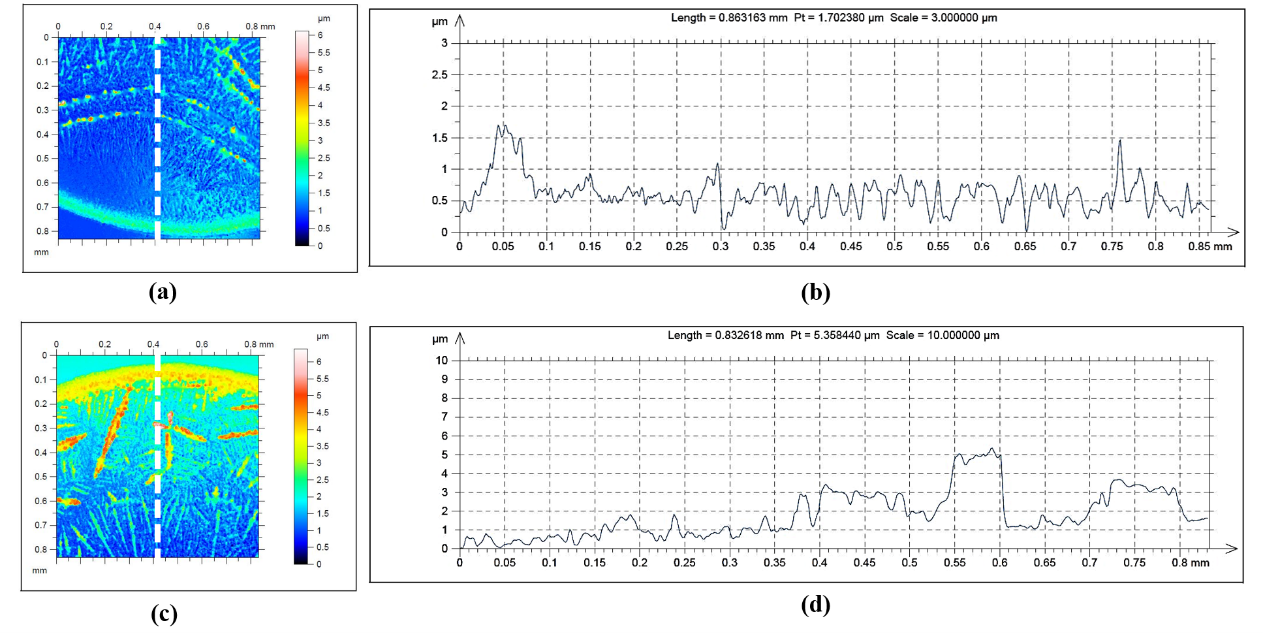

Figure S7 Optical profilometry data of dried precipitate for case 2, (a) 2D surface profile near side ‘1’, (b) Variation of the height along the dashed line in (a). (c) 2D surface profile near side ‘2’, (d) Variation of the height along the dashed line in (c).

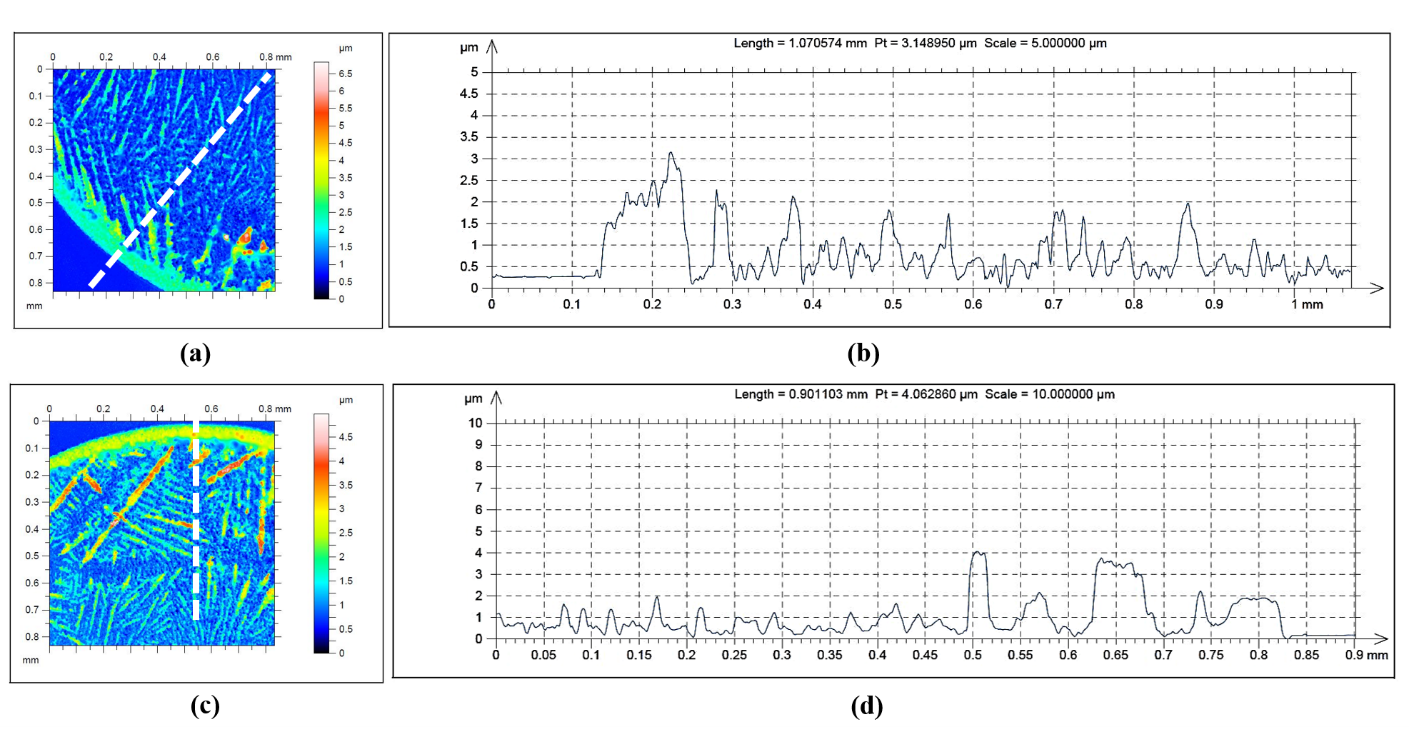

Figure S8 Optical profilometry data of dried precipitate for case 3, (a) 2D surface profile near side ‘1’, (b) Variation of the height along the dashed line in (a). (c) 2D surface profile near side ‘2’, (d) Variation of the height along the dashed line in (c).

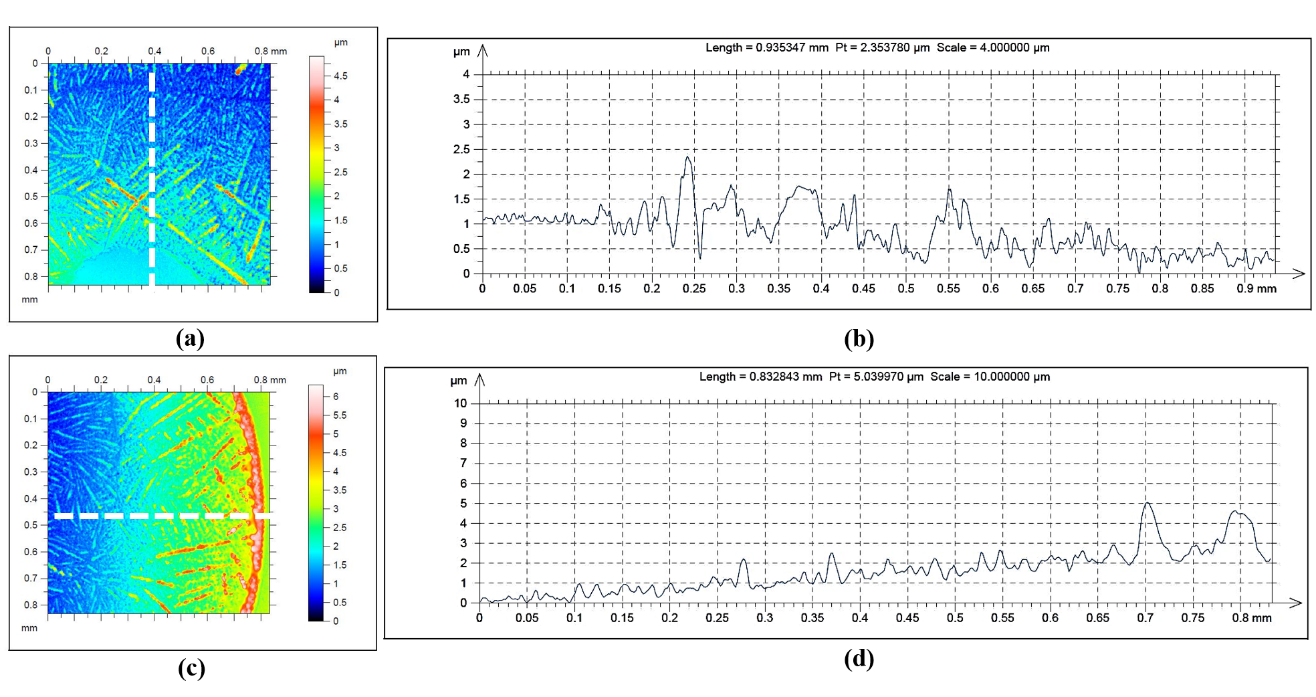

Figure S9 Optical profilometry data of dried precipitate for case 4, (a) 2D surface profile near side ‘1’, (b) Variation of the height along the dashed line in (a). (c) 2D surface profile near side ‘2’, (d) Variation of the height along the dashed line in (c).

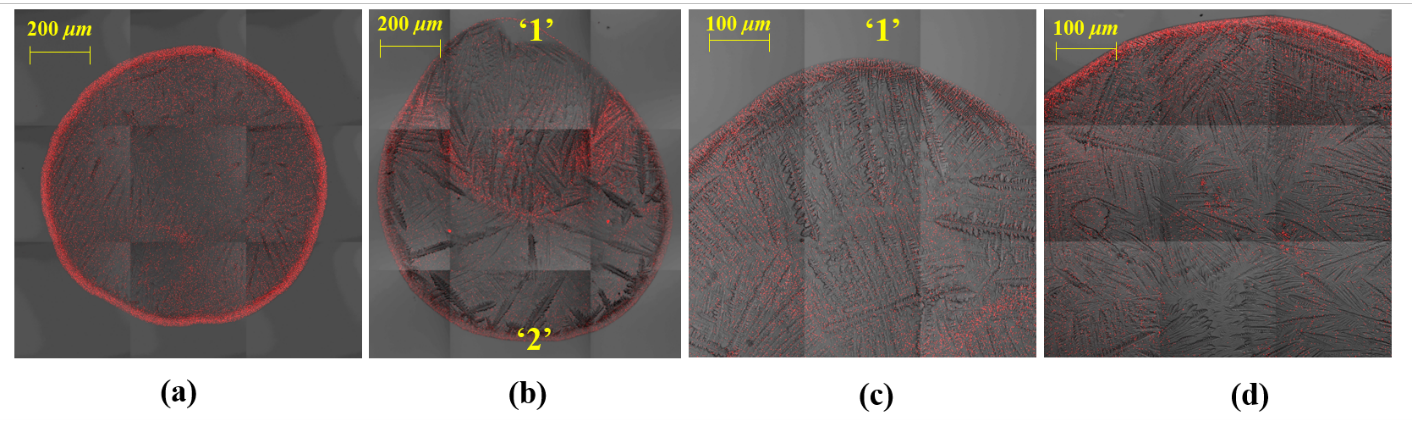

Figure S10 Confocal microscopy of the deposit with the fluorescence emission from the spherical polystyrene particles, (a) case 1, (b) case 2, (c) case 3, (d) case 4. Plots in the bottom row correspond to the intensity variation along the dashed line in the top row.

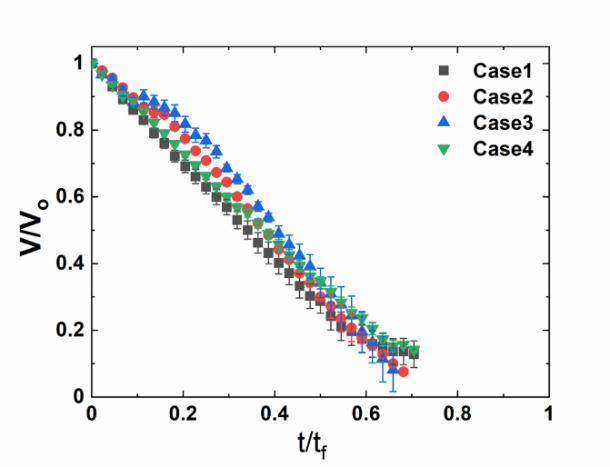

Figure S11 Volume regression curve of the SRF droplet for different cases.
